# Accurate ethnicity prediction from placental DNA methylation data

**DOI:** 10.1101/618470

**Authors:** Victor Yuan, E Magda Price, Giulia F Del Gobbo, Sara Mostafavi, Brian Cox, Alexandra M. Binder, Karin B. Michels, Carmen Marsit, Wendy P. Robinson

**Author notes:** **Corresponding Author**: Wendy Robinson.

## Abstract

**Background:** The influence of genetics on variation in DNA methylation (DNAme) is well documented. Yet confounding from population stratification is often unaccounted for in DNAme association studies. Existing approaches to address confounding by population stratification using DNAme data may not generalize to populations or tissues outside those in which they were developed. To aid future placental DNAme studies in assessing population stratification, we developed an ethnicity classifier, PlaNET (<u>Pla</u>cental DNAme Elastic <u>N</u>et <u>E</u>thnicity <u>T</u>ool), using five cohorts with Infinium Human Methylation 450k BeadChip array (HM450k) data from placental samples that is also compatible with the newer EPIC platform.

**Results:** Data from 509 placental samples was used to develop PlaNET and show that it accurately predicts (accuracy = 0.938, kappa = 0.823) major classes of self-reported ethnicity/race (African: n = 58, Asian: n = 53, Caucasian: n = 389), and produces ethnicity probabilities that are highly correlated with genetic ancestry inferred from genome-wide SNP arrays (>2.5 million SNP) and ancestry informative markers (n = 50 SNPs). PlaNET’s ethnicity classification relies on 1860 HM450K microarray sites, and over half of these were linked to nearby genetic polymorphisms (n = 955). Our placental-optimized method outperforms existing approaches in assessing population stratification in placental samples from individuals of Asian, African, and Caucasian ethnicities.

**Conclusion:** PlaNET provides an improved approach to address population stratification in placental DNAme association studies. The method can be applied to predict ethnicity as a discrete or continuous variable and will be especially useful when self-reported ethnicity information is missing and genotyping markers are unavailable. PlaNET is available as an R package at (https://github.com/wvictor14/planet).

## INTRODUCTION

Epigenome-wide association studies (EWAS) have shown that a substantial amount of variation in DNA methylation (DNAme) exists between human populations [1–7]. Therefore, if left unaccounted for, population-associated variation can interfere with the discovery of DNAme alterations associated with disease or environment. This type of confounding, often referred to as population stratification, can be addressed by inferring population-associated variation directly from DNAme data itself [8–10], as is done in genome-wide association studies (GWAS) [11]. However, unlike genetic markers, epigenetic markers are tissue-specific, and therefore a DNAme-based method developed in a specific tissue or population may not generalize well to other tissues with unique DNAme profiles.

In EWAS, confounding from population stratification is most often addressed using self-reported ethnicity/race to stratify study samples across the phenotype of interest. But, defining ethnicity/race is a complex task requiring the interpretation of a combination of biological and social factors leading to several complications: (i) inconsistent definition of ethnicity/race categories between individuals/organizations [12, 13]; (ii) self-reporting more than one ethnicity/race [14]; and (iii) missing ethnicity information altogether. To overcome the limitations of ethnicity/race categories, genetically-defined ancestry can be used [15] as an alternative measure of population-specific variation. In contrast to the discrete nature of ethnicity/race categories, genetic ancestry can be expressed as several continuous variables that reflect ancestry composition [16].

Though the use of genetic ancestry could help to better design EWAS, genotyping markers might not be collected in DNAme studies. In cases where self-reported ethnicity and genetic ancestry information are unavailable, methods have been developed to infer this information directly from DNAme (Table 1) measured on the popular Infinium Human Methylation 450k Beadchip array (HM450K) [8–10, 17]. Barfield et al. 2014 [8] and EPISTRUCTURE [9] methods both utilize principal components analysis (PCA) on select DNAme sites to infer genetic ancestry. Since only DNAme sites that are associated with nearby genetic variation are used, these methods produce principal components (PCs) that are often highly correlated with genome-wide genetic variation [8, 9], and therefore can be used as a measurement of genetic ancestry. Zhou et al. 2017 [10] explored using the set of 65 SNPs measured on HM450K to produce ethnicity/race classifications. However, it has not been investigated whether these methods perform well in populations and tissues other than the ones they were developed and tested in (Table 1).

**Table 1.** Description of methods to infer self-reported ethnicity or genetic ancestry using HM450K data.

| Name of method | Statistical Approach | Input HM450K sites | Output | Sample Characteristics |  |  |
| --- | --- | --- | --- | --- | --- | --- |
|  |  |  |  | Tissue | Populations* | Cohort Location |
| Barfield et al. 2014 [8] | PCA | 7703 DNAm sites with a 1000 genomes project SNP at the CpG site | Genetic ancestry as PC scores | Blood | Caucasian-Americans, African-Americans | USA |
| EPISTRUCTURE [9] | PCA | 4913 DNAm sites associated with local genetic variation (mQTLs) | Genetic ancestry as PC scores | Blood | Europeans, Puerto Ricans, Mexicans | Southern Germany; USA |
| Zhou et al. 2017 [10] | Predictive-modeling | 59/65 SNP sites | Ethnicity | Multiple | White, Black or African American, Asian | Many |
| PlaNET; this study | Predictive-modeling | 15 SNPs; 1845 DNAm sites | Ethnicity and Genetic Ancestry | Placenta | Caucasians, Asians, Africans | Canada, USA |

\* Ethnicity/ancestry as defined in associated study.

DNAme studies using placental tissue is of particular interest because the functioning of the placenta is essential to a healthy pregnancy [18, 19]. Although many DNAme alterations associated with placental-mediated diseases have been identified [20–23], the incidence of many of these conditions vary by population [24–26]. In this study we developed PlaNET (Placental DNAme Elastic Net Ethnicity Tool), an ethnicity classifier using DNAme and genotyping data measured on the HM450K array in multiple cohorts of placentas from North America. PlaNET was developed on overlapping sites from HM450K and the newer Illumina MethylationEPIC BeadChip array (EPIC) to ensure compatibility with future studies. We show that PlaNET out-performs existing methods in predicting ethnicity in placental tissue and can produce accurate measures of genetic ancestry. Importantly, our method can be used to classify individuals into discrete ancestral populations (i.e., African, Asian, and Caucasian) or to describe individuals on an ancestral continuum that may more accurately reflect the nature of modern human populations. In studies where ethnicity information is unavailable, PlaNET can be applied to predict ethnicity after obtaining DNAme data, and used to investigate population-specific differences or to minimize confounding by population stratification in statistical analyses.

## RESULTS

### Datasets

Our goal was to develop a placental DNAme-based ethnicity classifier, which could learn ethnicity-specific DNAme patterns from one set of samples in order to assign ethnicity labels to a new set of samples. We searched for placental HM450K data on the Gene Expression Omnibus [27] that contained more than one ethnicity group and made sample-specific ethnicity information available (Table 2). Five distinct cohorts met these criteria (labelled C1-C5), with three major North American ethnicities represented by sufficiently large numbers across more than one dataset: African (n = 58), Asian (n = 53), and Caucasian (n = 389). We opted to include samples from both healthy and abnormal pregnancies (preeclampsia, gestational diabetes mellitus, fetal growth restriction or overgrowth) (Table 2) [21, 28–33]. Though there were significant cohort-specific effects on DNAme that may reflect batch/technical variation (Additional file 2: Figure S1), we included these multiple datasets and phenotypes to enable the development of a robust classifier that would generalize well in future studies [34].

**Table 2.** Description of HM450K DNAme datasets used to develop and test PlaNET.

| Cohort<br>(n) | GEO<br>Accession | Dataset Summary | Location | Self-Reported Ethnicity* |  |  | Non-HM450K<br>Genetic<br>Data (n) |
| --- | --- | --- | --- | --- | --- | --- | --- |
|  |  |  |  | AFR<br>(n=57) | ASI<br>(n=53) | CAU<br>(n=389) |  |
| C1<br>(72) | GSE70453 | 36 controls, 36 gestational diabetes mellitus | Boston, MA, USA | 13 | 13 | 46 | N/A |
| C2<br>(24) | GSE73375 | 13 controls, 11 preeclampsia (PE) | Chapel Hill, NC, USA | 13 | 1 | 10 | N/A |
| C3<br>(289) | GSE75248 | 289 samples from infants with variable newborn neurobehaviour | RI, USA;<br>MA, USA | 23 | 9 | 257 | N/A |
| C4 | GSE100197 | 17 controls, 27 PE | Toronto, | 7 | 12 | 25 | 50 AIMs |
| (44) |  |  | CAN |  |  |  | (41) |
| C5<br>(70) | GSE100197,<br>GSE108567,<br>GSE74738,<br>Unpublished | 35 controls, 35<br>fetal growth<br>restriction, PE,<br>and/or preterm<br>birth | Vancouver,<br>CAN | 1 | 18 | 51 | 50 AIMs<br>(67);<br>Omni2.5<br>(27) |

\*AFR: African, ASI: Asian, CAU: Caucasian.

### Development of a placental DNA methylation ethnicity classifier

To determine the best machine learning classification algorithm that could learn ethnicity-specific patterns from DNAme microarray data, we compared four algorithms previously shown to be well-suited for prediction using high-dimensional genomics data [34–36]: generalized logistic regression with an elastic net penalty (GLMNET) [37, 38], nearest shrunken centroids (NSC) [35], k-nearest neighbours (KNN) [39], and support vector machines (SVM) [40]. For each algorithm, hyperparameter(s) were selected (e.g. k number of neighbours for KNN) that resulted in the highest performance estimated by repeated five-fold cross validation (three repeats). All algorithms performed favorably (logLoss = 0.170 - 0.276; Additional file 2: Figure S2a), except KNN (logLoss = 1.82). However, all algorithms showed a bias for high predicatability of Caucasians (average accuracy = 0.980), and low predictability of Asians (average accuracy = 0.448) (Additional file 2: Figure S2b). Considering overall- and ethnicity-specific performance, the GLMNET algorithm was used for the remainder of the study (accuracy = 0.866, 0.625, 0.998 for Africans, Asians, and Caucasians, respectively), and we refer to this classifier as PlaNET (<u>Pla</u>cental DNAme Elastic <u>N</u>et <u>E</u>thnicity <u>T</u>ool).

For each sample, PlaNET returns a probability that the sample is African, Asian or Caucasian and the final classification is defined by the ethnicity class with the highest of these probabilities. We reason that these probabilities have the potential to identify samples with mixed ancestry or ethnicity. Therefore, we implemented a threshold function on PlaNET’s probability outputs that classifies samples as ‘Ambiguous’ if the highest of the three class-specific probabilities is below 0.75 (Material and Methods, Additional file 2: Figure S3). This resulted in 7 self-reported African, 12 Asian, and 13 Caucasian samples as being classified as ambiguous, which led to a slight decrease in performance (Figure 1a). However, we note that because genetic ancestry is on a continuum and due to the limitations of self-reported ethnicity, there are likely to be individuals of mixed ancestry/ethnicity in our sample set, and therefore hypothesize that a model that includes an ambiguous class is more realistic and accurate than one without. Cross validation, where training/validation subsets were created based on cohort-identity, yielded an overall accuracy of 0.900, a Kappa of 0.738, and a positive predictive value of 0.944 (Figure 1a), which was consistent when examining performance by dataset (Additional file 2: Figure S4).

**Figure 1.**
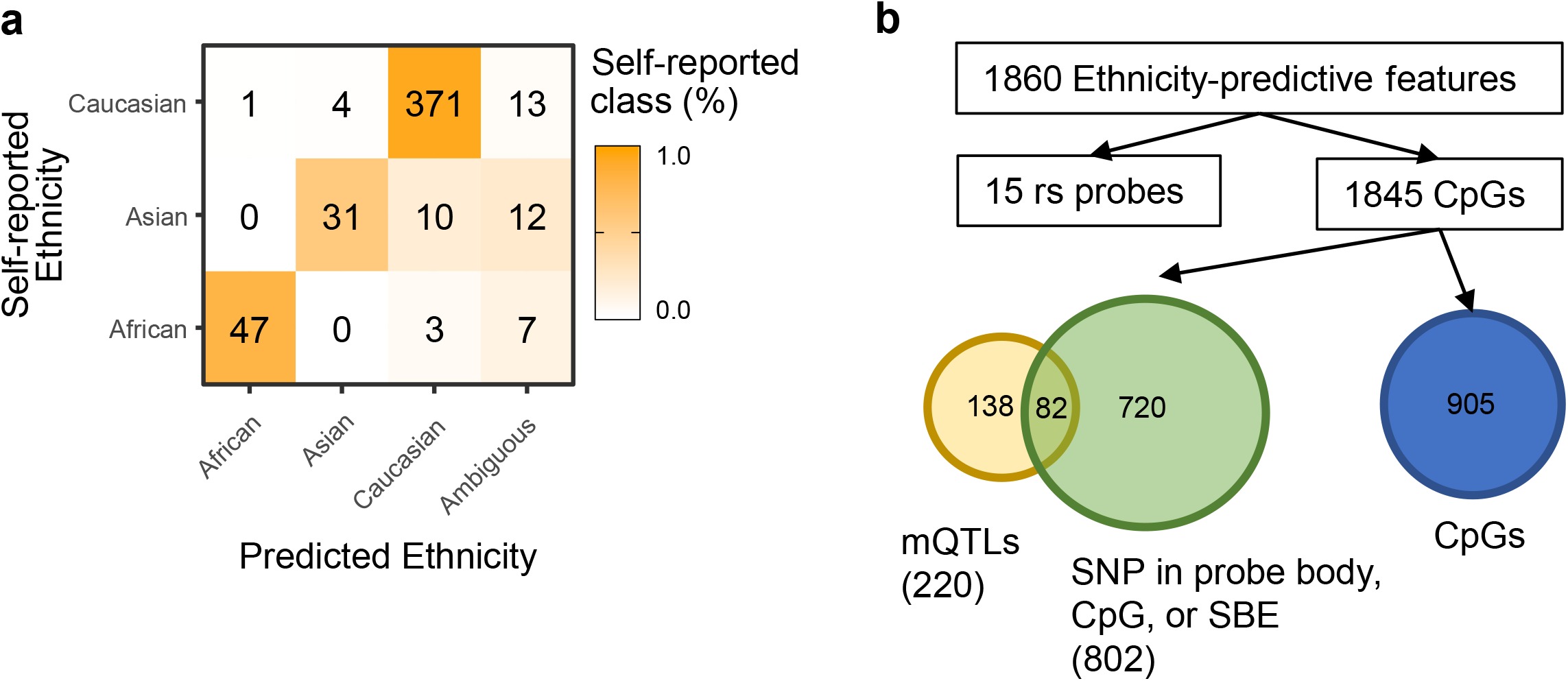
Evaluating PlaNET’s performance and characterizing ethnicity-predictive HM450K sites. We developed PlaNET (Placental elastic net ethnicity classifier), using placental HM450K data, and evaluated its classification performance using leave-one-dataset-out cross validation. **a** Each sample’s ethnicity classification from PlaNET is shown with respect to their self-reported ethnicity. Samples were called ‘ambiguous’ if their predicted probability fell below a ‘confidence’ threshold of 75%. **b** PlaNET utilizes a subset of ethnicity-predictive sites from the HM450K. To investigate whether genetic signal is present in the measurement for these sites, we cross-referenced ethnicity-predictive sites to an existing placental mQTL database [42] and determined whether any sites had SNPs present in either the probe body, CpG site of interrogation, or single base extension sites, based on dbSNP137.

### Ethnicity-predictive sites on the HM450K array are largely linked to genetic variation

To better understand the basis of PlaNET’s ethnicity prediction, we examined the 1860 sites [Additional file 1: Table S1] automatically-selected by the GLMNET model. These sites were enriched for SNP probes, containing 15 of the 59 SNPs explicitly measured on both HM450K and EPIC DNAme arrays (p < 1e-16). Of the remaining 1845 DNAme sites, we found significant enrichment for sites linked to genetic variation: 802 sites (43.1%) have a documented SNP in either the probe body, CpG site of interrogation, or the single base extension site (p < 1e-16) [41], and 220 sites (11.8%) corresponded to previously identified placental-specific methylation quantitative trait loci (mQTLs) [42] (p < 1e-16, Figure 1b). With respect to chromosomal location, we found significant enrichment for ethnicity-predictive sites on chromosomes 2 (p < 0.01), 15 (p < 0.05), and 17 (p < 0.05) (Additional file 2: Figure S5a). With respect to CpG density, we found significant enrichment for ethnicity-predictive sites in OpenSea (p < 0.001) and South Shore (p < 0.05) regions (Additional file 2: Figure S5b), where relatively neutral (unselected) genetic variation is more likely to be located [43]. Pathway analysis for GO and KEGG terms for genes associated with the 1860 sites, found only one significant (p < 0.05) GO term (homophilic cell adhesion via plasma membrane adhesion molecules).

### DNAme-inferred ethnicity and genetic ancestry

To test the ability of PlaNET to identify individuals of mixed ancestry, we examined whether samples classified as ‘ambiguous’ were also intermediate with respect to genetically-defined ancestry. Genetic ancestry was inferred from 50 ancestry informative genotyping markers (AIMs) in samples from cohorts C4 and C5 (n = 109), using 1000 Genomes Project samples as reference populations [44, 45]. These 50 markers were previously selected based on their ability to differentiate between African, European, East Asian, and South Asian populations [45]. Plotting the first two multi-dimensional scaling coordinates calculated on the 50 AIMs in (Figure 2), shows a handful of samples intermediate to three more distinct ancestry clusters. The samples with less extreme genetic ancestry coordinates based on AIMs tended to have lower PlaNET-calculated probabilities associated with the ethnicity classification matching the individual’s self-reported ethnicity (Figure 2), confirming that PlaNET provides some information on the genetic ancestry composition.

**Figure 2.**
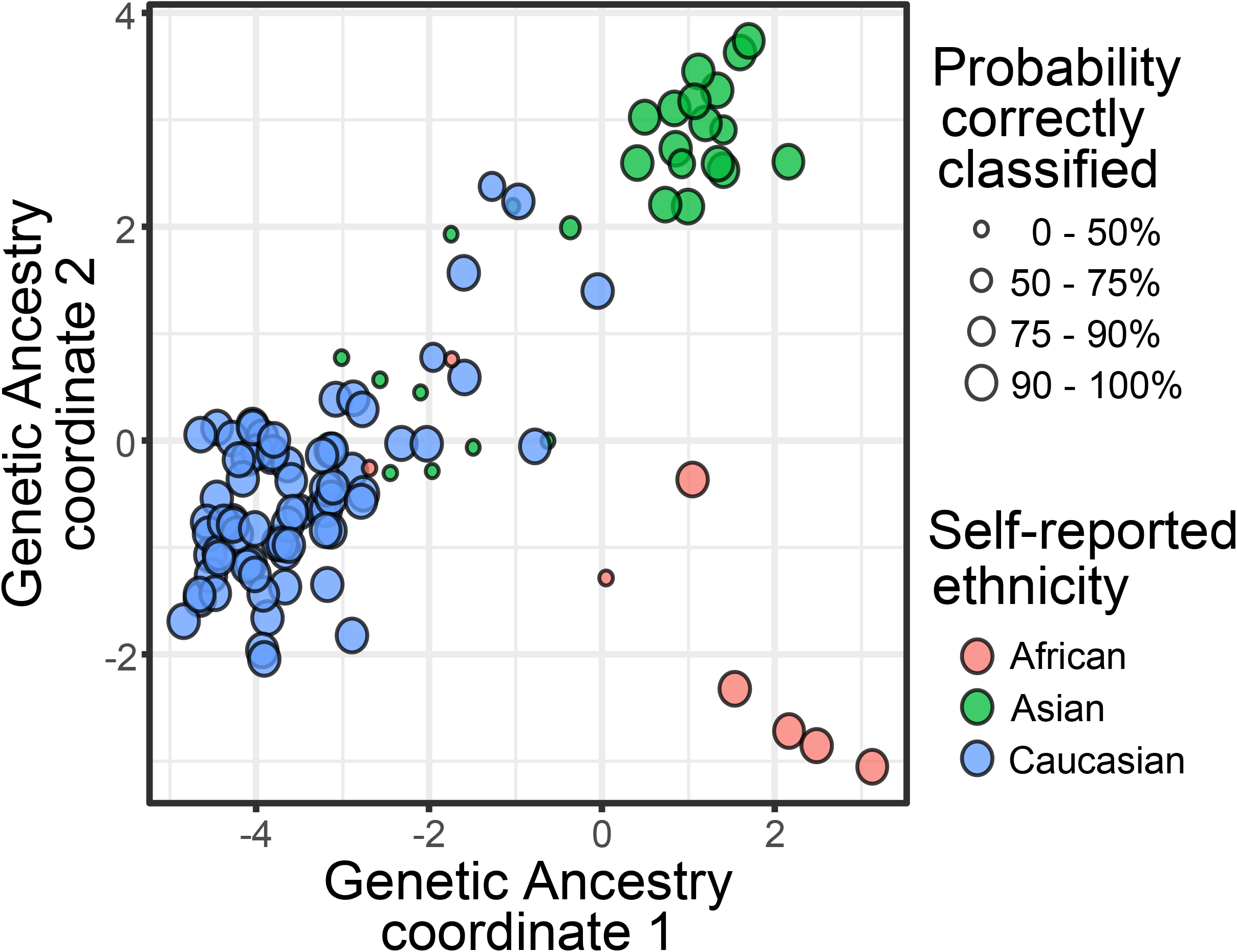
Probabilities associated with PlaNET ethnicity predictions and genetic ancestry inferred from AIMs. Ethnicity classifications from PlaNET and associated confidence/probability scores were compared to genetic ancestry inferred from 50 AIMs (n = 109, cohorts C4, C5), represented by the first three coordinates from multidimensional scaling using 1000 genomes project samples as reference populations.

Although genetic ancestry can be adequately inferred from a small set of AIMs, it is best obtained from a large number of unlinked markers [46]. Therefore, we also inferred genetic ancestry in a smaller number of samples from C5 (n = 37) with high density genotyping array data (Omni 2.5, >2.5 million SNPs), again using 1000 Genomes Project samples as reference populations [44, 47, 48], and compared this to PlaNET’s predicted membership probabilities for each ethnicity (Figure 3a-c). 10 of these 37 samples were not initially used for previous analyses due to a lack of available self-reported ethnicity information (Figure 3a). We found that genetic ancestry coefficients reflected the probabilities associated with ethnicity classification to a high degree (Figure 3bc, R^2^ = 0.95-0.96, p < 0.001).

**Figure 3.**
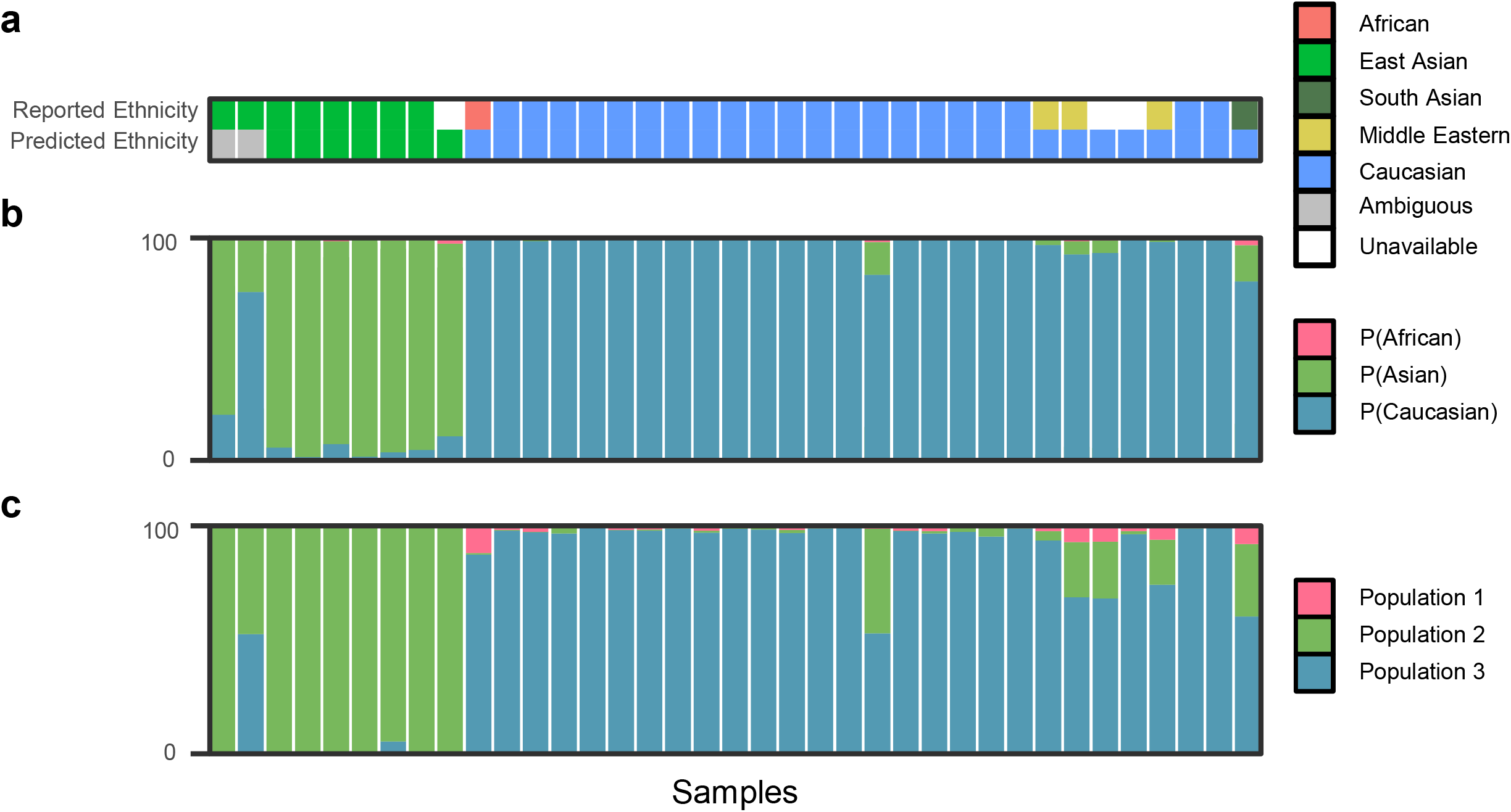
Probabilities associated with PlaNET ethnicity predictions and genetic ancestry inferred from high density genotyping data. PlaNET was tested in a subset of cohort C5 (n = 37). **a** PlaNET’s ethnicity classifications were compared with self-reported ethnicity. **b** Ethnicity probabilities generated by PlaNET were compared to c genetic ancestry coefficients determined from high density genotyping data (Omni 2.5, >2 million SNPs), using the function snmf() from the R package *LEA*, and found to be highly correlated (R^2^ = 0.95-0.96, p < 0.001) determined by linear regression.

### Characterizing existing methods to infer population structure in placental DNA methylation data

To evaluate our hypothesis that a placental-specific approach to population inference would outperform existing methods developed in other tissues, we compared the performance of PlaNET to three previously published HM450K methods: Barfield’s SNP-based filtering approach [8], EPISTRUCTURE [9], and Zhou’s SNP-based classifier [10]. To address the differences in the type of outcomes produced by each method (e.g. PCs or ethnicity classifications), we used PCA to generate metrics that could be compared between methods. PCA was performed on the set of HM450K sites corresponding to each method (Table 1) to determine the amount of variance explained in self-reported ethnicity (Figure 4a; n = 499, cohorts C1-C5), genetic ancestry (Figure 4b,c; n = 109, cohorts C4 and C5 only), and cohort-specific patient variables (e.g. microarray batch, sex, gestational age; Additional file 2: Figure S6), by each of the top ten PCs corresponding to each of the four population inference methods. For computation of PCs on PlaNET’s sites, we used a cohort-specific cross validation framework to account for bias that could be introduced by using the same samples for development and testing. Specifically, PlaNET’s PCs were computed separately for each cohort using ethnicity-predictive sites selected in all other cohorts (methods).

**Figure 4.**
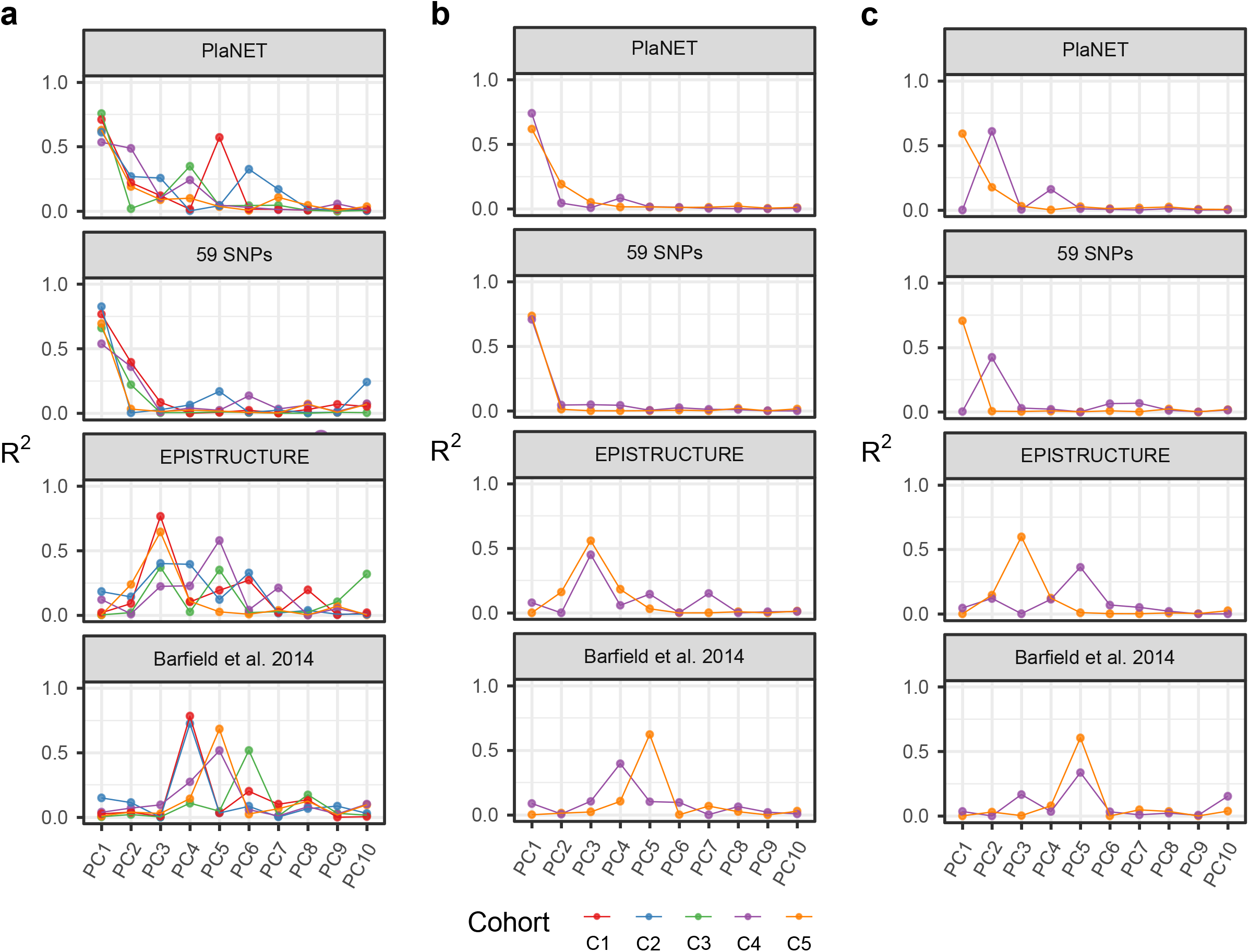
Comparing PlaNET to existing methods to account for population stratification using HM450K data. For each cohort, principal components analysis was conducted on PlaNET using a model trained on all other cohorts. PlaNET’s principal components (PCs) were then compared to the PCs computed on sites from EPISTRUCTURE [9], Barfield’s method [8], and the 59 SNPs. **a** Amount of variance explained in the first ten PCs for each method was calculated using a linear model with self-reported ethnicity as an independent variable. This was then repeated using AIMs in cohorts C4 and C5 for **b** genetic coordinate 1 and c genetic coordinate 2 (n = 109).

We found that for all cohorts, the first two PCs computed on PlaNET’s sites and the 59 SNPs was highly correlated with self-reported ethnicity (Figure 4a, R^2^ = 0.649 ± 0.087, 0.697 ± 0.110, respectively), genetic ancestry coordinate 1 (Figure 4b, R^2^ = 0.680 ± 0.086, 0.721 ± 0.019), and genetic ancestry coordinate 2 (Figure 4c, R^2^ = 0.296 ± 0.418, R^2^ = 0.356 ± 0.497; Figure 4a). In contrast, the first PC computed on Barfield’s and EPISTRUCTURE’s sites showed almost no correlation with self-reported ethnicity (Figure 4a, R^2^ = 0.0452 ± 0.060, 0.066 ± 0.082), genetic ancestry coordinate 1 (Figure 4b, R^2^ = 0.044 ± 0.060, 0.040 ± 0.055, respectively), or genetic ancestry coordinate 2 (Figure 4c, R^2^ = 0.0178 ± 0.0236, 0.0228 ± 0.0321). Instead, for Barfield and EPISTRUCTURE, the PCs that correlated with ethnicity/ancestry were confined to PCs 3-6 (Figure 4a), while often the top PCs (e.g., 1-4) for these two methods were associated with variables other than ethnicity/ancestry (Additional file 2: Figure S6). For example, in cohort C4, EPISTRUCTURE PC1 was most correlated with row position on the HM450K array (R^2^ = 0.482), PC2 with gestational age (R^2^ = 0.315), PC3 with genetic ancestry coordinate 1 (R^2^ = 0.450) and PC5 with ethnicity (R^2^ = 0.579; Additional file 2: Figure S6).

Limiting to methods that predict ethnicity classes, we compared the performance of PlaNET to Zhou et al. 2018’s SNP-based classifier (Additional file 2: Figure S7). Both classifiers demonstrated similar accuracy in classifying self-reported Africans (87.1% for PlaNET; 90.3% for Zhou) and Caucasians (96.7% vs 97.9%), but PlaNET was more accurate in classifying self-reported Asians (74.4% vs 41.0%).

### Application of PlaNET in an EWAS setting

Lastly, to demonstrate the utility of applying PlaNET to placental DNAme data, we applied PlaNET to obtain ethnicity classifications across two previously published EWAS studies using three datasets (Table 3, Additional file 2: Figure S9). We note that this includes samples from cohorts C4 and C5 that were used to develop PlaNET.

**Table 3.**
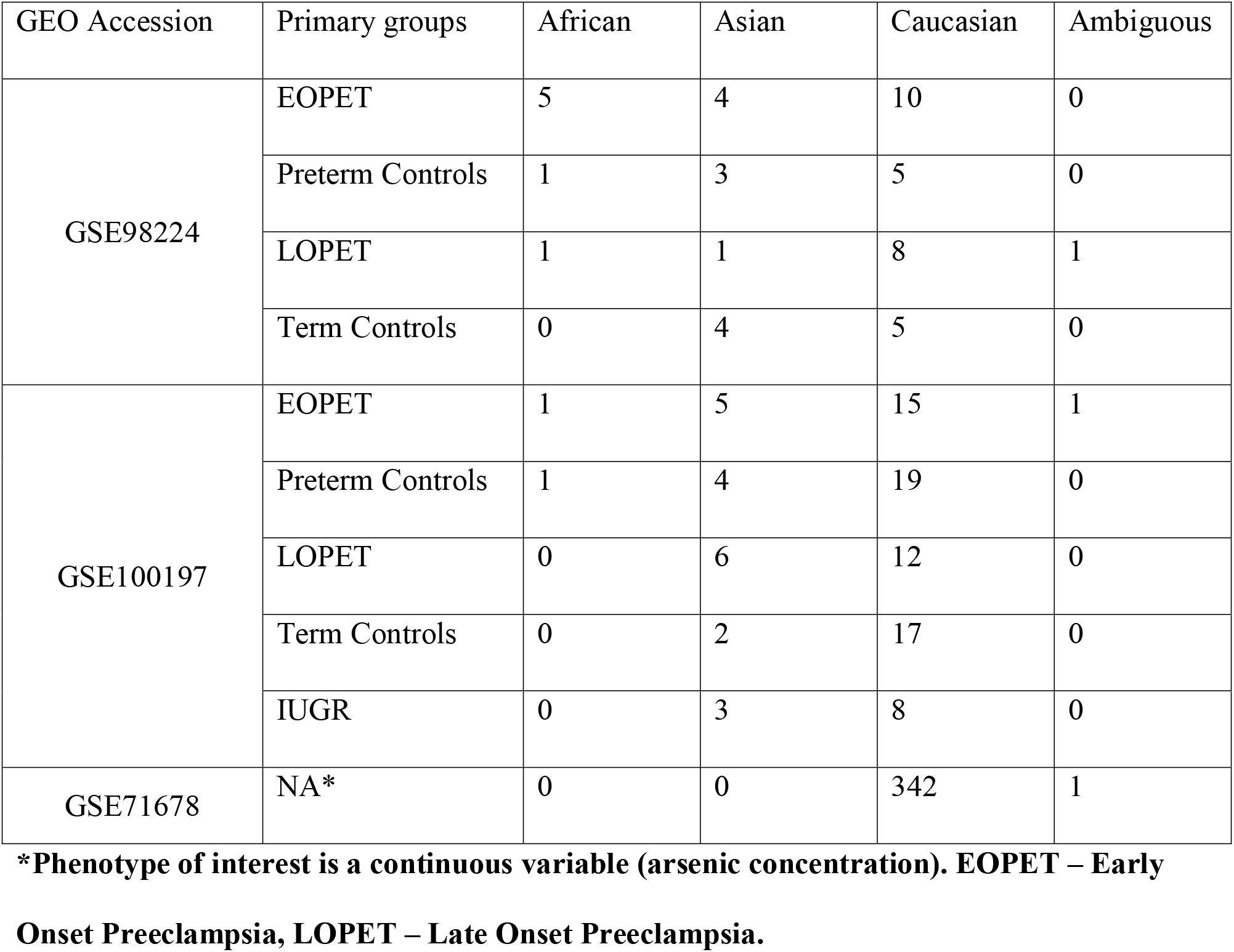
Distribution of PlaNET ethnicity predictions across previously published placental EWAS datasets.

One study used discovery GSE100197 (n = 102) and validation GSE98224 (n = 48) datasets to investigate DNAme alterations associated with preeclampsia status [21]. We reasoned that correction for ethnicity should decrease false positives in the EWAS and therefore increase concordance between hits identified in the two data sets. In the original EWAS, with no adjustment for ethnicity, our group reported that 599 out of the 1703 (35.1%) significant associations found in the discovery cohort were also significant in the validation cohort, and the correlation of the difference in mean DNAme between controls and preeclampsia-affected samples (i.e. delta betas) at FDR significant sites between discovery and validation was 0.62 [21]. When we repeated the analysis while adjusting for ethnicity determined by PlaNET, the number of preeclampsia-associated sites that overlapped between cohorts increased to 651/1614 (40.3%) [Additional file 1: Table S5], and the correlation between delta betas increased to 0.66. We also found that repeating gene set enrichment analysis, which originally found nothing significant [21], yielded several significantly enriched GO terms such as developmental process, inflammatory response and cell adhesion [Additional file 1: Table S6]. Next, because adjustment for population stratification can not only be done via correction in linear modelling, but can also be done by stratifying an analysis by population identity, we performed a secondary EWAS confined to samples predicted as Caucasians (n = 71/102 for discovery, n = 28/48 for validation). This resulted in a decrease in overlap in preeclampsia-associated sites between cohorts: 359/1488 (17%) [Additional file 1: Table S7], although the correlation between delta betas remained high (r = 0.67), indicating the observed decrease in overlap between significantly differentially methylated sites was likely due to a decrease in power from smaller sample size (particularly in the validation group) rather than a decrease in concordance between cohorts.

PlaNET can be useful for checking for discrepancies in self-reported ethnicity information. We tested whether PlaNET could identify the ethnicity of samples from an all-Caucasian population. GSE71678 (n = 343), a cohort not used in the development of PlaNET, consisted of DNAme data from placental samples collected from a New Hampshire, USA birth cohort that investigated the effects of arsenic exposure on placental DNAme [49]. PlaNET determined 342 samples were classified as Caucasian, and 1 sample had a high probability of belonging to the Caucasian group (Probability = 0.73) but was below our confidence threshold and was therefore classified as ‘ambiguous’, confirming ethnic homogeneity was high in this cohort and adjustment for population stratification was not needed in this study.

## DISCUSSION

In this study, we developed PlaNET, a method to predict Asian, African, and Caucasian ethnicity using placental HM450K array data. To enable compatibility with future studies, PlaNET was developed on sites (452,453 CpGs and 59 SNPs) overlapping between HM450K and EPIC Illumina DNAme arrays. Although all samples in this study were reported as a single ethnicity/race, we expected that there would be significant population substructure that might limit our ability to develop predictive models of ethnicity and to assess their performance. Despite this limitation, ethnicity could be predicted with high accuracy as assessed by cross validation. PlaNET’s DNAme-based ethnicity classification relies on HM450K sites with large amounts of genetic signal, which supported our initial efforts to filter our data to enrich for genetic-informative sites prior to classifier development (methods) [41, 50, 51]. When examining PlaNET’s 1,860 sites used to predict ethnicity, more than half could be linked to a nearby genetic polymorphism. Of these, 802 CpG sites have documented SNPs in their probe body, single base extension or CpG site of interrogation, which previously have been identified to differ between European and East Asian populations [41]. Several studies have suggested the genetic influence on DNAme at these sites is primarily technical in nature [41, 50, 51], suggesting the patterns in DNAme at these sites are likely tissue-agnostic, warranting further investigation in their utility in predicting ethnicity and/or genetic ancestry in tissues other than the placenta. A significant proportion of other ethnicity-predictive CpG sites (n = 220) were previously found associated with placental mQTLs in a population with similar demographics to the ones studied here [42]. This finding, together with EPISTRUCTURE—a method that also relies on mQTLs [9]—suggests that leveraging the tissue- and population-specificity of mQTLs can produce highly effective DNAme-based population structure inference methods.

Of the existing methods to assess population stratification from DNAme data, we note that Barfield’s method and EPISTRUCTURE infer continuous measures of genetic ancestry, while Zhou’s SNP-based classifier returns discrete ethnicity classifications, however ours produce both [8–10] (Table 1). EPISTRUCTURE and Barfield’s method are unsupervised PCA-based approaches, which rely on the empirical observation that specific DNAme sites can be highly correlated with PCs computed on genome-wide genotype data in adult blood samples [8, 9]. However, we found that DNAme at these sites did not produce PCs that are highly associated with genotype data in placental samples. Instead, top PCs were more often associated with non-ancestry related variables in the placental samples included in this study, such as gestational age, preeclampsia, and technical variables. Ethnicity and genetic ancestry-associated PCs were confined to the third to sixth component of variation, suggesting that application of these methods may require filtering of PCs to those that are ethnicity / ancestry-specific, which is impossible when self-reported ethnicity and genetic ancestry information is unavailable (i.e. when these methods are needed most). Future improvements to these types of methods can aim at improving the amount of ethnicity and genetic ancestry-associated signal in the sites used to ensure the top two-three PCs are always associated with ethnicity and ancestry. This aim could also be supported in identifying ethnicity and ancestry-associated sites that are also robust to changes in non-genetic drivers of DNAme such as cell type, gestational age, and severe pathology.

Supervised population inference approaches such as ethnicity classifiers can return an explicit assignment of samples into distinct ancestral groups. In comparison to self-reported ethnicity, an assessment based on DNAme/genetic data is more objectively defined, which allows for more robust investigation of ethnicity-specific effects. An important goal of any population structure inference method would be to identify samples of mixed ancestry, a capability not well supported by Zhou’s ethnicity classifier [10]. In contrast, PlaNET produced membership probabilities corresponding to each ethnicity group that were highly correlated with genetic ancestry estimated from genotyping data. This was consistent whether we used principal components analysis on AIMs data, or model-based estimation of ancestry on high density genotyping array data [47, 52–54]. In this study, we defined samples of potential mixed ancestry as those with a maximum membership probability of less than 0.75, but we note that this threshold can be manually adjusted by the user, and that the probabilities themselves can be used to adjust for population structure in study populations including significant numbers of samples with mixed ancestry.

Results of DNAme studies on genetic ancestry and ethnicity, such as this one, depend on the number and proportion of different populations sampled from, as well as the tissue studied. Due to limitations in sample availability, only African, Asian, and Caucasian ethnicities were included in our study. However, we note that these ethnicities are among the most common in North American populations—but future developments should consider inclusion of additional ethnicities. Furthermore, due to limited number of samples with high density genetic data, we were unable to address the extent of finer population structure that likely exists within the major ancestral groups studied. Differences in ethnic composition in samples from our study and samples used to develop Barfield’s method and EPISTRUCTURE may also explain why Barfield’s method or EPISTRUCTURE performed poorly in our study [8, 9]. A lack of generalizability of these methods to our placental samples was likely further compounded by the use of different tissues to develop each method—Barfield and EPISTRUCTURE were both developed and tested in blood tissue only. This is especially important to consider when applying these techniques to tissues with unique DNAme profiles, such as placenta [18]. It is possible that application of these approaches to other tissues that are more similar to blood (e.g. other somatically-derived tissues) may result in better performance compared to when applied to placenta as seen in this study. However, any DNAme-based test needs to be validated before application to new tissues, which has not yet been done for these methods.

A major goal of EWAS is to uncover signal truly associated with the phenotype/environment of interest that might generalize to other relevant populations. This is challenging given the wide host of technical variables that can affect DNAme measurements and the common finding that many phenotypes are associated with relatively small effect sizes [33, 55]. To this end, adjustment for major confounders such as genetic ancestry or ethnicity can significantly improve EWAS. We demonstrated, in a reanalysis of our previously published PE placentas, that adjustment for ethnicity, determined by PlaNET, improved the replicability of significant associations between independent cohorts. Conversely, overadjustment can occur when populations are relatively homogeneous, resulting in bias and/or loss of precision. We showed that PlaNET can indicate minimal population stratification when applied to a homogenous Caucasian population. Thus, PlaNET will be useful in assessing population stratification in future placental EWAS, as well as conducting ethnicity-stratified analyses, which may lead to important insights into the disparities between populations of pregnancy-related outcomes [24–26].

## CONCLUSIONS

We demonstrated that ethnicity and genetic ancestry can be accurately predicted using placental HM40K DNAme microarray data with respect to three major ethnicity/ancestral populations. Although samples that were used to develop PlaNET were reported to come from single ethnic populations, our classifier was able to capture mixed ancestry, and outperformed existing prediction methods. PlaNET will be valuable in assessing and accounting for population stratification, which can confound associations between DNAme with disease or environment, in future studies using HM450K or EPIC arrays. The machine-learning approach used to develop PlaNET can easily be applied for other tissues and populations for use in future DNAme studies.

## METHODS

### Collection of previously published placental HM450K DNA methylation data

Placental DNAme data from liveborn deliveries of healthy and mixed pregnancy complications (n = 585), were combined from seven GEO HM450K datasets corresponding to five North American cohorts (summarized in Table 2; sample-specific information in Additional file 1: Table S4) [21, 27, 29–32]. Five unpublished samples from the C5 cohort were included and are available at GSE128827. Gestational ages of these pregnancies at delivery ranged from 26 to 42 weeks and 50.30% of samples were male. Samples were excluded (n=67) if their self-reported ethnicity was missing or did not fall into one of three major race/ethnicity groups: Asian/East Asian (n=53), Caucasian/White (non-hispanic) (n=389), or African/African American/Black (n=57). Based on census data [56], we note that self-reported Caucasian/White (non-hispanic) samples are typically of European ancestry, self-reported Asians are typically of East Asian ancestry and self-reported Africans represent diverse ancestries from Africa with a significant potential of admixture from other ancestries [57]. When possible, data was downloaded as raw IDAT files (GSE75248, GSE100197, GSE100197, GSE108567, GSE74738), otherwise methylated and unmethylated intensities were utilized (GSE70453, GSE73375).

### DNA methylation data processing

All samples were analyzed using the Illumina Infinium HumanMethylation450 BeadChip array (HM450K), the most popular measure of DNAme for EWAS. Array data analysis was performed using R version 3.5.0. To allow compatibility of PlaNET with the newest Infinium MethylationEPIC BeadChip array (EPIC), the raw HM450K data (485,512 CpGs, 65 SNPs) was filtered to the 452,453 CpGs and 59 SNPs common between both platforms prior to classifier development [10]. Because genetic variability can capture ancestry information, we omitted the common filtering step that would remove sites with probes that overlap SNPs (n = 52,116 at a minor allele frequency > 0.05). CpGs were removed if greater than 1% of samples had poor quality signal (bead count < 3, or a detection p-value >0.01; n = 14,858). The remaining poor quality measurements were replaced with imputed values using K-Nearest Neighbours from the R package *impute* [58]. Cross-hybridizing (n = 41,937) [50, 51] and placental-specific non-variable sites (n = 86,502) [59] were also removed, leaving 319,233 sites for classifier development.

Biological sex was determined by hierarchical clustering on DNAme measured from sites on the sex chromosomes and then compared to reported sex. Samples with discordant reported and inferred sex were removed (n=3). Samples were also removed if they had a low mean inter-array correlation (< 0.95, n = 5). Intra-array normalization methods, normal-exponential out-of-band (NOOB) [60] and beta mixture quantile normalization (BMIQ) [61] were used from R packages *minfi* (version 1.26.2) [62] and *wateRmelon* (version 1.24.0) [63] to normalize data.

### Genotyping data collection and genetic ancestry assessment

In a subset of C5 (n = 27) and 10 additional samples, high density SNP array genotypes were collected. DNA samples from one site from the fetal side of each placenta were collected as previously described [45] and quality was checked using a NanoDrop ND-1000 (Thermo Scientific) as well as by electrophoresis on a 1% agarose gel. Genotyping at ~2.3 million SNPs was done on the Illumina Infinium Omni2.5-8 (Omni2.5) array at the Centre for Applied Genomics, Hospital for Sick Kids, Toronto, Canada. For inferring genetic ancestry, the data for these 37 samples was combined with a previously processed 1000 Genomes Project Omni2.5 dataset (n = 1,756) to use as reference populations [44, 48]. Genotypes in this combined dataset were filtered for quality (missing call rate > 0.05, n removed = 31,604), minor allele frequency (MAF > 0.05, n removed = 114,628), and linkage disequilibrium pruning was performed to select representative SNPs (R^2^ < 0.25, n removed = 919,824) for a final dataset of 218,732 SNPs and n = 1793 samples. Genetic ancestry coefficients were estimated using the R package *LEA*, which utilizes sparse non-negative matrix factorization to produce similar results to model-based algorithms ADMIXTURE and STRUCTURE [47, 54]. Cross-entropy criterion was used to assess the number of ancestral populations (Additional file 2: Figure S8) [64].

A smaller panel of 50 ancestry-informative genotyping markers (AIMs) was collected in a subset of samples from cohorts C4 (n = 41) and C5 (n = 68). AIMs were selected based on their ability to differentiate between African, European, East Asian, and South Asian populations [65–67]. Results from cohort C5 have been published elsewhere [45], and genotyping data was collected for cohort C4 in the same manner. Briefly, these markers were measured in placental villus DNA using the Sequenom iPlex Gold platform (Génome Québec Innovation Centre, Montréal, Canada). Genetic ancestry inferred from 50 AIMs markers was computed using multi-dimensional scaling after combining with the same 50 AIMs from the 1000 Genomes Project samples, as previously described [45].

### Developing the ethnicity classifier and assessing its performance

To develop and assess the performance of PlaNET we used a ‘leave-one-dataset-out cross-validation’ (LODOCV) approach. This approach uses four out of five datasets to develop a predictive model (training), which is then used to generate ethnicity classifications on the samples in the remaining dataset (testing). This differs from the traditional cross validation approach of randomly splitting the full dataset into training and testing. LODOCV produces more accurate estimates of classifier performance for future studies, and has been previously used for evaluating age-predictive models [34]. Each iteration of LODOCV generates dataset-specific estimates of performance (accuracy, Kappa). After all iterations, overall performance was assessed by aggregating classifications across all datasets.

For fitting predictive models within LODOCV-generated training sets, we used the R package *caret* [68]. Several algorithms were compared: logistic regression with an elastic net penalty (GLMNET) [37, 38], nearest shrunken centroids (NSC) [35, 69], K-nearest neighbours (KNN) [39], and support vector machines (SVM) [40]. To determine optimum tuning parameters for each algorithm (e.g., ‘k’ number of neighbours for KNN, alpha and lambda for GLMNET), we built several models while varying the tuning parameter(s) and compared the performance of these models within each training set using repeated (n = 3) five-fold cross validation. Hyperparameter values were left as default settings in *caret* [68], or a grid of values for GLMNET (alpha = 0.025 – 0.500, lambda = 0.0025 – 0.2500). We compared the performance of these models using accuracy, positive predictive value, cohen’s Kappa [70], and logLoss (a measure of classification accuracy that heavily penalizes over-confident misclassifications). The results from this analysis can be found in Additional file 1: Table S2, S3. After assessing the classifier performance using LODOCV, a final GLMNET model was fit to the entire dataset (cohorts C1-C5) using the same model fitting procedure described above and is available for use in future datasets (https://github.com/wvictor14/planet).

### Enrichment analysis

The DNAme sites [see Additional file 1: Table S1] and SNPs selected to predict ethnicity in this final model (n = 1860) were used for enrichment analysis. For DNAme sites, we looked for enrichment for SNPs in the probe body, CpG site, and single base extension sites based on Illumina’s HM450K annotation version 1.2 [71]. We looked for enrichment for placental mQTLs [42], chromosomes and CpG islands (HG19; Additional file 2: Figure S5). Fisher’s exact test was used for all enrichment tests using a p-value threshold of < 0.05, and was carried out in R using the function fisher.test(). GO and KEGG pathway analysis was done using the R package *missMethyl* version 3.8 [72].

### Threshold analysis

We explored the use of a ‘threshold function’ to identify samples that are difficult to classify into discrete ethnicity groupings because of mixed ancestry. Because PlaNET’s ethnicity classifications are associated with varying degrees of confidence (i.e., probabilities), we reasoned that a sample’s most probable ethnicity classification (i.e., max(P(Asian), P(African), P(Caucasian)) would be lower with a higher degree of mixed ancestry. Therefore, we implemented a threshold function on PlaNET’s probability outputs that classifies samples as ‘Ambiguous’ if the highest of the three class-specific probabilities is below a certain threshold. We explored several thresholds and decided on 0.75, which minimized the resulting decrease in predictive performance (Additional file 2: Figure S3).

### Comparison of methods for inferring genetic ancestry / ethnicity from HM450K data

Because existing population inference methods and PlaNET use different statistical approaches to infer genetic ancestry/ethnicity (PCA-based vs predictive modeling), we compared each method based on the amount of population-associated signal in DNAme from each method-specific subset of sites. This was done by applying principal component analysis (PCA) to standardized beta values for HM450k sites associated with each method (Table 1) [8–10] within each cohort. To avoid bias, the PCs associated with PlaNET were calculated for each cohort using a classifier trained on all other cohorts (generated from LODOCV).

To test for the amount ethnicity and genetic ancestry –associated signal in the sites corresponding to each method, we applied several simple linear regression models to estimate the amount of variance explained in PC_i_ (i = 1, 2, 3, …, 10) by self-reported ethnicity and genetic ancestry when available. To determine other factors that might affect signal in these sites, we also tested for the association between PCi and each covariate available for each cohort. All simple regression tests were done in R using the function lm().

To compare PlaNET to Zhou et al. 2017’s SNP-based classifier [10], we used the package R package *sesame* (version 1.1.0) [73] to obtain SNP-based ethnicity classifications for samples with idats available (cohorts C3, C4, and C5).

### Application of PlaNET to previous EWAS

To demonstrate application of PlaNET, we downloaded placental HM450K DNAme datasets GSE98224, GSE100197, and GSE71678. We note that GSE100197 and GSE98224 overlap cohorts C4 and C5, respectively. To apply PlaNET to obtain ethnicity information, raw data was downloaded from GEO in the form of IDATs and loaded into R using *minfi* (version 1.26.2). Both NOOB and BMIQ normalization were applied before applying PlaNET. The R package *limma* (version 3.36.2) was used to test for differentially methylated sites. For GSE98224 and GSE100197, the processed DNAme data was used, and statistical thresholds were chosen the same as the published analysis [21]. For enrichment analysis, differentially methylated CpGs were inputted into the *gometh* function from the R package *missMethyl* (version 1.16.0) using all filtered sites as background, and default settings.

## Abbreviations

PlaNET: Placental DNAme Elastic Net Ethnicity Tool
DNAme: DNA methylation
CpG: Cytosine-phosphate-guanine
SNP: Single-nucleotide polymorphism
AIMs: Ancestry informative genotyping markers
mQTL: methylation quantitative trait loci
PCA: Principal component analysis
PC: Principal component
HM450K: Infinium HumanMethylation450 BeadChip
EPIC: Infinium MethylationEPIC BeadChip
LODOCV: Leave-one-dataset-out cross validation
GLMNET: Generalized logistic regression with an elastic net penalty
SVM: Support vector machines
KNN: K-nearest neighbours
NSC: Nearest shrunken centroids
PlaNET: Placental elastic net ethnicity classifier
USA: United States of America
AFR: African
ASI: Asian
CAU: Caucasian
BMIQ: Beta-mixture interquantile normalization
NOOB: Normal exponential out-of-band normalization.

## DECLARATIONS

### Ethics approval and consent to participate

The bulk of the data used in this study is from public repositories. Additional molecular investigations of the Vancouver cohort is covered under Ethics approval obtained from the University of British Columbia and BC Women’s and Children’s Hospital research ethics board in Vancouver, BC, Canada (certificate numbers H16-02280, H04–704488).

### Consent for publication

Not applicable.

### Availability of data and material

PlaNET can be downloaded as an R package from (https://github.com/wvictor14/planet). For reproducibility, all source code for development of software and analysis in this study is available at (https://github.com/wvictor14/Ethnicity_Inference_450k) and deposited on Zenodo (doi: https://doi.org/10.5281/zenodo.2641272 for source code, doi: https://doi.org/10.5281/zenodo.2641266 for software) [74, 75].

The processed DNAme data supporting the conclusions of this article are available in the Gene Expression Omnibus repository (https://www.ncbi.nlm.nih.gov/geo/query/acc.cgi?acc=GSE128827). Raw DNAme data for previously published samples is available in the repositories listed in Table 2. Raw DNAme data for samples published for the first time used in this study are available also at GSE128827. Genetic ancestry coordinates are available in supporting documents along with other sample-specific information (Additional file 1: Table S4). The genotyping array data used to generate genetic ancestry coordinates are not publicly available due to privacy concerns.

### Authors’ contributions

VY, EMP, GDG, SM, and WR contributed to the design of the study. VY, EMP, and GDG performed all data analysis. EMP, GDG, SM, BC, AMB, KBM, CM, WR helped generate and contribute the data. EMP and WR conceived of the study. All authors approved and contributed to the writing of the final manuscript.

## Acknowledgements

Advice and support from Louie Dinh, Dr. Lisa McEwan, Dr. Sam Wilson, Chaini Konwar, Amy Inkster, and Dr. Maria Penaherrera are greatly appreciated. Special thanks to Dr. Catherine Fry for correspondance regarding GEO data.

## Additional materials

| File name | File format | Title of data | Descriptive title | Description of data |
| --- | --- | --- | --- | --- |
| Additional file 1 | .xls | Table S1 | Ethnicity-predictive HM450K sites | A list of all PlaNET's 1860 automatically-selected ethnicity-predictive HM450K sites. |
| Additional file 1 | .xls | Table S2 | PlaNET's cohort-specific ethnicity classification performance. | PlaNET's classification performance described across cohorts, assessed using LODOCV. |
| Additional file 1 | .xls | Table S3 | PlaNET's ethnicity classification performance described by class. | PlaNET's classification performance described across ethnicity groups, assessed using LODOCV. |
| Additional file 1 | .xls | Table S4 | Sample metadata | A table of sample-specific information. |
| Additional file 1 | .xls | Table S5 | PE linear modeling results while adjusting for predicted ethnicity | Results containing 651 PE-associated sites from linear modeling while adjusting for predicted ethnicity. |
| Additional file 1 | .xls | Table S6 | PE linear modelling results in Caucasian cohort | Results containing 359 PE-associated sites from linear modeling in Caucasian-only samples. |
| Additional file 1 | .xls | Table S7 | GO enrichment analysis results | GO terms specifically enriched based on the 651 PE-associated sites obtained from linear modeling. |
| Additional file 2 | .pdf | Figure S1 | Dataset-specific effects | PC1 by PC2 scatterplot from PCA computed on scaled and centered DNAm beta values from 499 samples and 319233 sites. |
| Additional file 2 | .pdf | Figure S2 | Performance between machine learning algorithms in training | Resampling results for each machine learning algorithm. **a** performance (LogLoss) between machine learning algorithms in predicting ethnicity, and **b** class-specific accuracy. |
| Additional file 2 | .pdf | Figure S3 | Threshold analysis for determining "ambiguous" samples | Various cutoffs for predicted membership probabilities were compared with respect to changes in predictive performance. |
| Additional file 2 | .pdf | Figure S4 | Dataset-specific performance | PlaNET's classification performance was calculated for each dataset using a model trained to all other datasets. |
| Additional file 2 | .pdf | Figure S5 | Enrichment analysis on ethnicity-predictive HM450K sites | PlaNET's CpG sites used to predict ethnicity was tested for enrichment with respect to **a** chromosomal location, and **b** relation to CpG islands. |
| Additional file 2 | .pdf | Figure S6 | Association of population structure PCs with technical and biological variables | PCs were computed on ethnicity-predictive sites, EPISTRUCTURE, Barfield's method and the 59 SNP probes. Each PC was tested for their association with various cohort-specific technical and biological variables. For a given cohort (e.g. C1), ethnicity predictive sites from a classifier trained on all other cohorts (e.g. C2-C5) was used to avoid bias. |
| Additional file 2 | .pdf | Figure S7 | PlaNET vs Zhou et al. 2017 snp-based classifier | PlaNET's ethnicity classification performance was compared to Zhou et al. 2018 [76] SNP-based ethnicity classifier in cohorts C3, C4, and C5. |
| Additional file 2 | .pdf | Figure S8 | Estimating k number of ancestral populations using in genetic admixture inference program LEA | The cross-entropy criterion was used to determine the number of ancestral populations for estimating genetic ancestry coefficients. The number of ancestral populations was chosen at the point $k = 3$, when the cross-entropy criterion decreases significantly less with each integer-increase in $k$. |
| Additional file 2 | .pdf | Figure S9 | Application of PlaNET to placental EWAS | Samples from three independent cohorts are plotted along three axes by their probability of belonging to each ethnicity class and colored by their final ethnicity classification determined by PlaNET. |

## REFERENCES

1. Chu S-K, Yang H-C. Interethnic DNA methylation difference and its implications in pharmacoepigenetics. Epigenomics. 2017;9:1437–54. doi:10.2217/epi-2017-0046.

2. Smith JA, Zhao W, Wang X, Ratliff SM, Mukherjee B, Kardia SLR, et al. Neighborhood characteristics influence DNA methylation of genes involved in stress response and inflammation: The Multi-Ethnic Study of Atherosclerosis. Epigenetics. 2017;12:662–73. doi:10.1080/15592294.2017.1341026.

3. Park SL, Patel YM, Loo LWM, Mullen DJ, Offringa IA, Maunakea A, et al. Association of internal smoking dose with blood DNA methylation in three racial/ethnic populations. Clin Epigenetics. 2018;10:110. doi:10.1186/s13148-018-0543-7.

4. Adkins RM, Krushkal J, Tylavsky FA, Thomas F. Racial differences in gene-specific DNA methylation levels are present at birth. Birth Defects Res A Clin Mol Teratol. 2011;91:728–36. doi:10.1002/bdra.20770.

5. Kwabi-Addo B, Wang S, Chung W, Jelinek J, Patierno SR, Wang B-D, et al. Identification of differentially methylated genes in normal prostate tissues from African American and Caucasian men. Clin Cancer Res. 2010;16:3539–47. doi:10.1158/1078-0432.CCR-09-3342.

6. Liu J, Hutchison K, Perrone-Bizzozero N, Morgan M, Sui J, Calhoun V. Identification of genetic and epigenetic marks involved in population structure. PLoS One. 2010;5:e13209. doi:10.1371/journal.pone.0013209.

7. Fraser HB, Lam LL, Neumann SM, Kobor MS. Population-specificity of human DNA methylation. Genome Biol. 2012;13:R8. doi:10.1186/gb-2012-13-2-r8.

8. Barfield RT, Almli LM, Kilaru V, Smith AK, Mercer KB, Duncan R, et al. Accounting for Population Stratifcation in DNA methylation studies. 2014;38:231–41.

9. Rahmani E, Shenhav L, Schweiger R, Yousefi P, Huen K, Eskenazi B, et al. Genome-wide methylation data mirror ancestry information. Epigenetics Chromatin. 2017;10:1. doi:10.1186/s13072-016-0108-y.

10. Zhou W, Laird PW, Shen H. Comprehensive characterization, annotation and innovative use of Infinium DNA methylation BeadChip probes. Nucleic Acids Res. 2017;45:gkw967. doi:10.1093/nar/gkw967.

11. Tian C, Gregersen PK, Seldin MF. Accounting for ancestry: population substructure and genome-wide association studies. Hum Mol Genet. 2008;17:R143–50. doi:10.1093/hmg/ddn268.

12. Sankar P, Cho MK, Mountain J. Race and ethnicity in genetic research. Am J Med Genet A. 2007;143A:961–70. doi:10.1002/ajmg.a.31575.

13. Mersha TB, Abebe T. Self-reported race/ethnicity in the age of genomic research: its potential impact on understanding health disparities. Hum Genomics. 2015;9:1. doi:10.1186/s40246-014-0023-x.

14. Kaneshiro B, Geling O, Gellert K, Millar L. The challenges of collecting data on race and ethnicity in a diverse, multiethnic state. Hawaii Med J. 2011;70:168–71.

15. Banda Y, Kvale MN, Hoffmann TJ, Hesselson SE, Ranatunga D, Tang H, et al. Characterizing Race/Ethnicity and Genetic Ancestry for 100,000 Subjects in the Genetic Epidemiology Research on Adult Health and Aging (GERA) Cohort. Genetics. 2015;200:1285–95. doi:10.1534/genetics.115.178616.

16. Pritchard JK, Rosenberg NA. Use of Unlinked Genetic Markers to Detect Population Stratification in Association Studies. Am J Hum Genet. 1999;65:220–8.

17. Bibikova M, Barnes B, Tsan C, Ho V, Klotzle B, Le JM, et al. High density DNA methylation array with single CpG site resolution. Genomics. 2011;98:288–95. doi:10.1016/J.YGENO.2011.07.007.

18. Robinson WP, Price EM. The human placental methylome. Cold Spring Harb Perspect Med. 2015;5:a023044. doi:10.1101/cshperspect.a023044.

19. Robinson WP, Peñaherrera MS, Konwar C, Yuan V, Wilson SL. Epigenetic Modifications in the Human Placenta. Hum Reprod Prenat Genet. 2019;:293–311. doi:10.1016/B978-0-12-813570-9.00013-9.

20. Yeung KR, Chiu CL, Pidsley R, Makris A, Hennessy A, Lind JM. DNA methylation profiles in preeclampsia and healthy control placentas. Am J Physiol - Hear Circ Physiol. 2016;310:H1295–303. doi:10.1152/ajpheart.00958.2015.

21. Wilson SL, Leavey K, Cox BJ, Robinson WP. Mining DNA methylation alterations towards a classification of placental pathologies. Hum Mol Genet. 2018;27:135–46. doi:10.1093/hmg/ddx391.

22. Rong C, Cui X, Chen J, Qian Y, Jia R, Hu Y. DNA Methylation Profiles in Placenta and Its Association with Gestational Diabetes Mellitus. Exp Clin Endocrinol Diabetes. 2015;123:282–8. doi:10.1055/s-0034-1398666.

23. Konwar C, Price EM, Wang LQ, Wilson SL, Terry J, Robinson WP. DNA methylation profiling of acute chorioamnionitis-associated placentas and fetal membranes: insights into epigenetic variation in spontaneous preterm births. Epigenetics Chromatin. 2018;11:63. doi:10.1186/s13072-018-0234-9.

24. Bryant AS, Worjoloh A, Caughey AB, Washington AE. Racial/ethnic disparities in obstetric outcomes and care: prevalence and determinants. Am J Obstet Gynecol. 2010;202:335–43. doi:10.1016/j.ajog.2009.10.864.

25. Xiao J, Shen F, Xue Q, Chen G, Zeng K, Stone P, et al. Is ethnicity a risk factor for developing preeclampsia? An analysis of the prevalence of preeclampsia in China. J Hum Hypertens. 2014;28:694–8. doi:10.1038/jhh.2013.148.

26. Rosenberg TJ, Garbers S, Lipkind H, Chiasson MA. Maternal obesity and diabetes as risk factors for adverse pregnancy outcomes: differences among 4 racial/ethnic groups. Am J Public Health. 2005;95:1545–51. doi:10.2105/AJPH.2005.065680.

27. Edgar R, Domrachev M, Lash AE. Gene Expression Omnibus: NCBI gene expression and hybridization array data repository. Nucleic Acids Res. 2002;30:207–10. doi:10.1093/nar/30.1.207.

28. Leavey K, Wilson SL, Bainbridge SA, Robinson WP, Cox BJ. Epigenetic regulation of placental gene expression in transcriptional subtypes of preeclampsia. Clin Epigenetics. 2018;10:28. doi:10.1186/s13148-018-0463-6.

29. Martin E, Ray PD, Smeester L, Grace MR, Boggess K, Fry RC. Epigenetics and Preeclampsia: Defining Functional Epimutations in the Preeclamptic Placenta Related to the TGF-β Pathway. PLoS One. 2015;10:e0141294. doi:10.1371/journal.pone.0141294.

30. Paquette AG, Houseman EA, Green BB, Lesseur C, Armstrong DA, Lester B, et al. Regions of variable DNA methylation in human placenta associated with newborn neurobehavior. Epigenetics. 2016;11:603–13. doi:10.1080/15592294.2016.1195534.

31. Binder AM, LaRocca J, Lesseur C, Marsit CJ, Michels KB. Epigenome-wide and transcriptome-wide analyses reveal gestational diabetes is associated with alterations in the human leukocyte antigen complex. Clin Epigenetics. 2015;7:79. doi:10.1186/s13148-015-0116-y.

32. Hanna CW, Peñaherrera MS, Saadeh H, Andrews S, McFadden DE, Kelsey G, et al. Pervasive polymorphic imprinted methylation in the human placenta. Genome Res. 2016;26:756–67.

33. Price EM, Robinson WP. Adjusting for Batch Effects in DNA Methylation Microarray Data, a Lesson Learned. Front Genet. 2018;9:83. doi:10.3389/fgene.2018.00083.

34. Horvath S. DNA methylation age of human tissues and cell types. Genome Biol. 2013;14:R115. doi:10.1186/gb-2013-14-10-r115.

35. Tibshirani R, Hastie T, Narasimhan B, Chu G. Class Prediction by Nearest Shrunken Centroids, with Applications to DNA Microarrays. Stat Sci. 2003;18:104–17.

36. Huang S, Cai N, Pacheco PP, Narrandes S, Wang Y, Xu W. Applications of Support Vector Machine (SVM) Learning in Cancer Genomics. Cancer Genomics Proteomics. 2018;15:41–51. doi:10.21873/cgp.20063.

37. Friedman J, Hastie T, Tibshirani R. Regularization Paths for Generalized Linear Models via Coordinate Descent. J Stat Softw. 2010;33:1–22.

38. Zou H, Hastie T. Regularization and variable selection via the elastic net. J R Stat Soc B. 2005;67:301–20.

39. Altman NS. An Introduction to Kernel and Nearest-Neighbor Nonparametric Regression. Am Stat. 1992;46:175–85. doi:10.1080/00031305.1992.10475879.

40. Cortes C, Vapnik V. Support-vector networks. Mach Learn. 1995;20:273–97. doi:10.1007/BF00994018.

41. Daca-Roszak P, Pfeifer A, Żebracka-Gala J, Rusinek D, Szybińska A, Jarząb B, et al. Impact of SNPs on methylation readouts by Illumina Infinium HumanMethylation450 BeadChip Array: implications for comparative population studies. BMC Genomics. 2015;16:1003. doi:10.1186/s12864-015-2202-0.

42. Delahaye F, Do C, Kong Y, Ashkar R, Salas M, Tycko B, et al. Genetic variants influence on the placenta regulatory landscape. PLOS Genet. 2018;14:e1007785. doi:10.1371/journal.pgen.1007785.

43. Walser J-C, Furano A V. The mutational spectrum of non-CpG DNA varies with CpG content. Genome Res. 2010;20:875–82. doi:10.1101/gr.103283.109.

44. Gibbs RA, Boerwinkle E, Doddapaneni H, Han Y, Korchina V, Kovar C, et al. A global reference for human genetic variation. Nature. 2015;526:68–74. doi:10.1038/nature15393.

45. Del Gobbo GF, Price EM, Hanna CW, Robinson WP. No evidence for association of MTHFR 677C>T and 1298A>C variants with placental DNA methylation. Clin Epigenetics. 2018;10:34. doi:10.1186/s13148-018-0468-1.

46. Price AL, Zaitlen NA, Reich D, Patterson N. New approaches to population stratification in genome-wide association studies. Nat Rev Genet. 2010;11:459–63. doi:10.1038/nrg2813.

47. Frichot E, François O. LEA : An R package for landscape and ecological association studies. Methods Ecol Evol. 2015;6:925–9. doi:10.1111/2041-210X.12382.

48. Roslin NM, Li W, Paterson AD, Strug LJ. Quality control analysis of the 1000 Genomes Project Omni2.5 genotypes. 2016. doi:10.1101/078600.

49. Green BB, Karagas MR, Punshon T, Jackson BP, Robbins DJ, Houseman EA, et al. Epigenome-wide assessment of DNA methylation in the placenta and arsenic exposure in the New Hampshire Birth Cohort Study (USA). Environ Health Perspect. 2016;124:1253–60.

50. Chen Y, Lemire M, Choufani S, Butcher DT, Grafodatskaya D, Zanke BW, et al. Discovery of cross-reactive probes and polymorphic CpGs in the Illumina Infinium HumanMethylation450 microarray. Epigenetics. 2013;8:203–9. doi:10.4161/epi.23470.

51. Price ME, Cotton AM, Lam LL, Farré P, Emberly E, Brown CJ, et al. Additional annotation enhances potential for biologically-relevant analysis of the Illumina Infinium HumanMethylation450 BeadChip array. Epigenetics Chromatin. 2013;6:4. doi:10.1186/1756-8935-6-4.

52. Price AL, Patterson NJ, Plenge RM, Weinblatt ME, Shadick NA, Reich D. Principal components analysis corrects for stratification in genome-wide association studies. Nat Genet. 2006;38:904–9. doi:10.1038/ng1847.

53. Pritchard JK, Stephens M, Donnelly P. Inference of population structure using multilocus genotype data. Genetics. 2000;155:945–59.

54. Alexander DH, Novembre J, Lange K. Fast model-based estimation of ancestry in unrelated individuals. Genome Res. 2009;19:1655–64. doi:10.1101/gr.094052.109.

55. Breton C V, Marsit CJ, Faustman E, Nadeau K, Goodrich JM, Dolinoy DC, et al. Small-Magnitude Effect Sizes in Epigenetic End Points are Important in Children’s Environmental Health Studies: The Children’s Environmental Health and Disease Prevention Research Center’s Epigenetics Working Group. Environ Health Perspect. 2017;125:511–26. doi:10.1289/EHP595.

56. United States Census Bureau: Population and Housing. 2018. https://www.census.gov/en.html. Accessed 18 Jun 2018.

57. Bryc K, Durand EY, Macpherson JM, Reich D, Mountain JL. The genetic ancestry of African Americans, Latinos, and European Americans across the United States. Am J Hum Genet. 2015;96:37–53. doi:10.1016/j.ajhg.2014.11.010.

58. Hastie T, Tibshirani R, Narasimhan B, Chu G. impute: Imputation for microarray data. Bioconductor. 2018. http://bioconductor.org/packages/release/bioc/html/impute.html.

59. Edgar RD, Jones MJ, Robinson WP, Kobor MS. An empirically driven data reduction method on the human 450K methylation array to remove tissue specific non-variable CpGs. Clin Epigenetics. 2017;9:11. doi:10.1186/s13148-017-0320-z.

60. Triche TJ, Weisenberger DJ, Van Den Berg D, Laird PW, Siegmund KD, Siegmund KD. Low-level processing of Illumina Infinium DNA Methylation BeadArrays. Nucleic Acids Res. 2013;41:e90. doi:10.1093/nar/gkt090.

61. Teschendorff AE, Marabita F, Lechner M, Bartlett T, Tegner J, Gomez-Cabrero D, et al. A beta-mixture quantile normalization method for correcting probe design bias in Illumina Infinium 450 k DNA methylation data. Bioinformatics. 2013;29:189–96. doi:10.1093/bioinformatics/bts680.

62. Aryee MJ, Jaffe AE, Corrada-Bravo H, Ladd-Acosta C, Feinberg AP, Hansen KD, et al. Minfi: a flexible and comprehensive Bioconductor package for the analysis of Infinium DNA methylation microarrays. Bioinformatics. 2014;30:1363–9. doi:10.1093/bioinformatics/btu049.

63. Pidsley R, Y Wong CC, Volta M, Lunnon K, Mill J, Schalkwyk LC. A data-driven approach to preprocessing Illumina 450K methylation array data. BMC Genomics. 2013;14:293. doi:10.1186/1471-2164-14-293.

64. Frichot E, Mathieu F, Trouillon T, Bouchard G, François O. Fast and efficient estimation of individual ancestry coefficients. Genetics. 2014;196:973–83. doi:10.1534/genetics.113.160572.

65. Phillips C, Salas A, Sánchez JJ, Fondevila M, Gómez-Tato A, Álvarez-Dios J, et al. Inferring ancestral origin using a single multiplex assay of ancestry-informative marker SNPs. Forensic Sci Int Genet. 2007;1:273–80. doi:10.1016/J.FSIGEN.2007.06.008.

66. Phillips C, Freire Aradas A, Kriegel AK, Fondevila M, Bulbul O, Santos C, et al. Eurasiaplex: a forensic SNP assay for differentiating European and South Asian ancestries. Forensic Sci Int Genet. 2013;7:359–66. doi:10.1016/j.fsigen.2013.02.010.

67. Fondevila M, Phillips C, Santos C, Freire Aradas A, Vallone PM, Butler JM, et al. Revision of the SNPforID 34-plex forensic ancestry test: Assay enhancements, standard reference sample genotypes and extended population studies. Forensic Sci Int Genet. 2013;7:63–74. doi:10.1016/j.fsigen.2012.06.007.

68. Kuhn M. Building Predictive Models in *R* Using the caret Package. Journal of Statistical Software. 2008;28:1–26. doi:10.18637/jss.v028.i05.

69. Tibshirani R, Hastie T, Narasimhan B, Chu G. Diagnosis of multiple cancer types by shrunken centroids of gene expression. Proc Natl Acad Sci U S A. 2002;99:6567–72. doi:10.1073/pnas.082099299.

70. McHugh ML. Interrater reliability: the kappa statistic. Biochem medica. 2012;22:276–82.

71. Hansen KD. IlluminaHumanMethylation450kanno.ilmn12.hg19: Annotation for Illumina’s 450k methylation arrays. Bioconductor. 2016.

72. Phipson B, Maksimovic J, Oshlack A. missMethyl: an R package for analyzing data from Illumina’s HumanMethylation450 platform. Bioinformatics. 2015;32:btv560. doi:10.1093/bioinformatics/btv560.

73. Zhou W, Triche TJ, Laird PW, Shen H. SeSAMe: reducing artifactual detection of DNA methylation by Infinium BeadChips in genomic deletions. Nucleic Acids Res. 2018;46:e123–e123. doi:10.1093/nar/gky691.

74. Yuan V, Price EM, Del Gobbo G, Mostafavi S, Cox B, Binder AM, et al. wvictor14/PlaNET: PlaNET. Zenodo. 2019. doi:10.5281/zenodo.2641266.

75. Yuan V, Price EM, Del Gobbo G, Mostafavi S, Cox B, Binder AM, et al. wvictor14/Ethnicity_Inference_450k: analysis scripts. Zenodo. 2019. doi:10.5281/zenodo.2641272.

76. Qi Y-H, Teng F, Zhou Q, Liu Y-X, Wu J-F, Yu S-S, et al. Unmethylated-maspin DNA in maternal plasma is associated with severe preeclampsia. Acta Obstet Gynecol Scand. 2015;94:983–8. doi:10.1111/aogs.12691.

