## Additional file 2 for "Accurate ethnicity prediction from placental DNA methylation data"

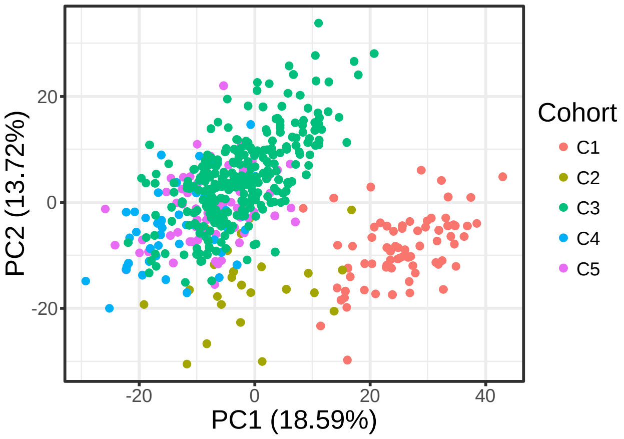


**Figure S1. Dataset-specific effects.** PC1 by PC2 scatterplot from PCA computed on scaled and centered DNAme beta values from 499 samples and 319233 sites.

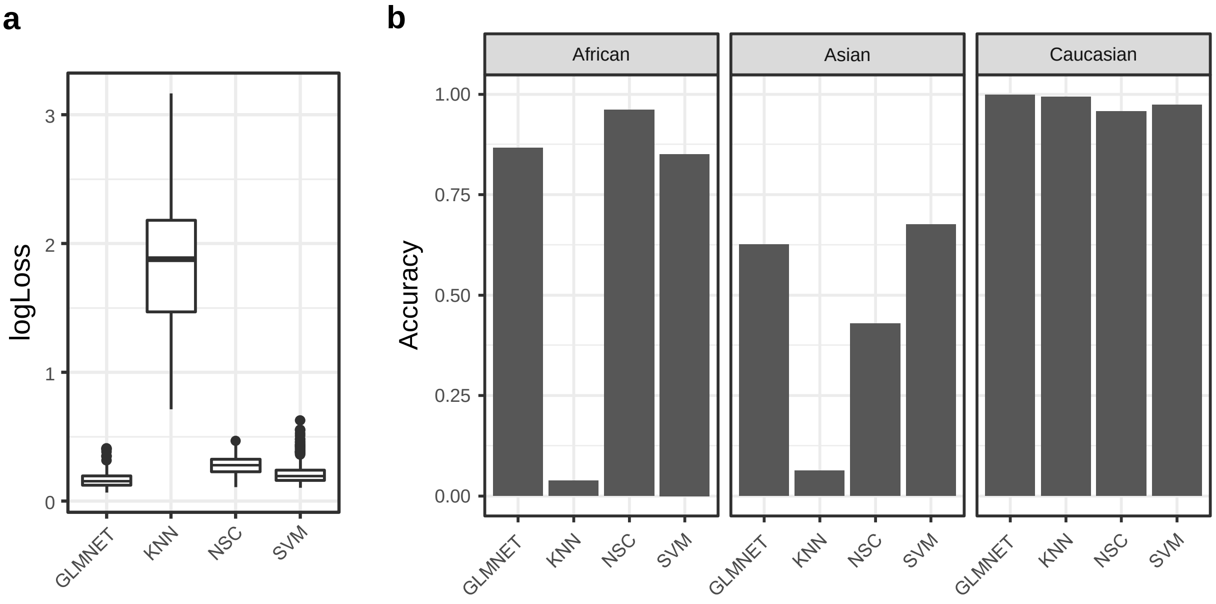


**Figure S2. Performance between machine learning algorithms in training.** Resampling results for each machine learning algorithm. **a** performance (LogLoss) between machine learning algorithms in predicting ethnicity, and **b** class-specific accuracy.

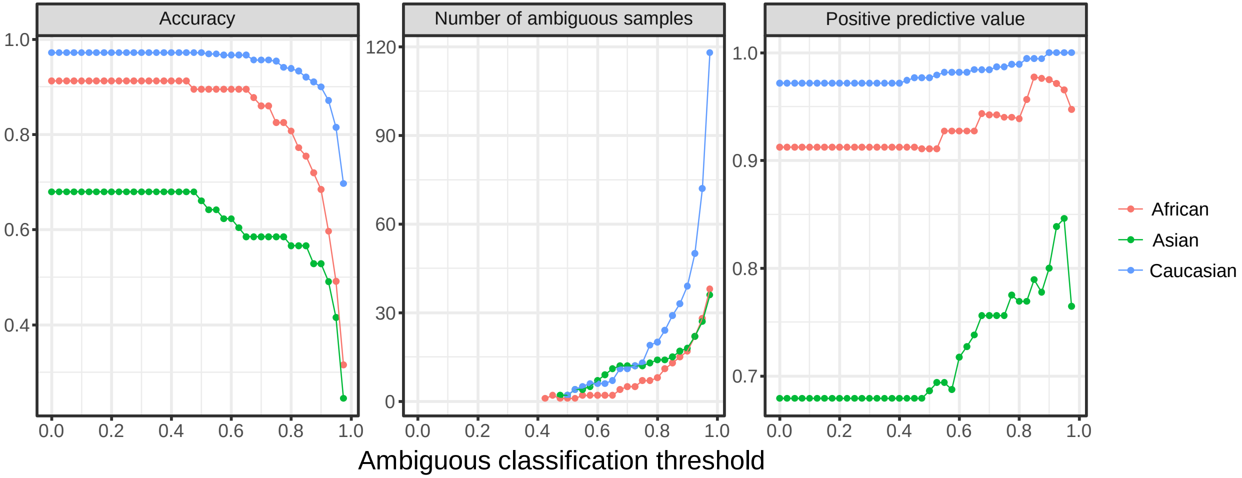


**Figure S3. Threshold analysis for determining "ambiguous" samples.** Various cutoffs for predicted membership probabilities were compared with respect to changes in predictive performance.

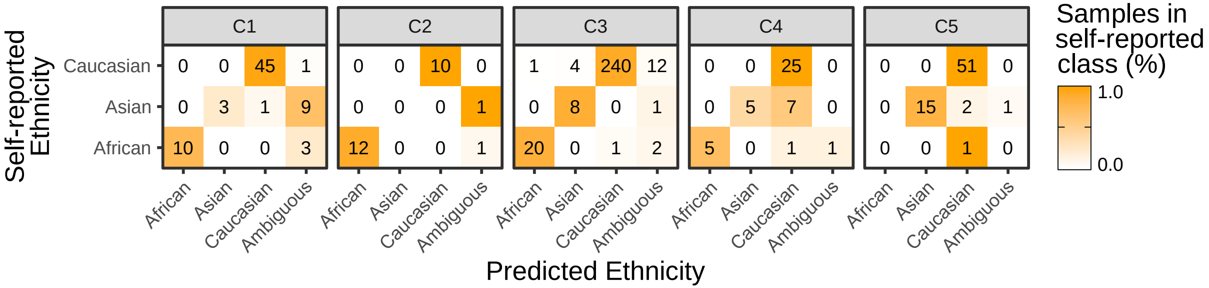


**Figure S4. Dataset-specific performance.** PlaNET's classification performance was calculated for each dataset using a model trained to all other datasets.

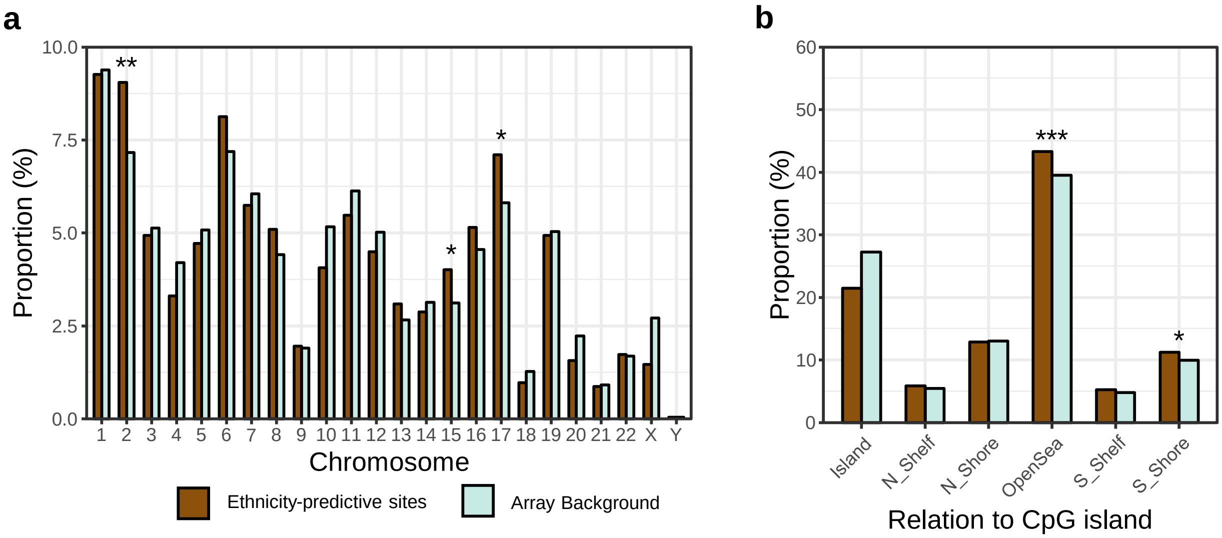


**Figure S5. Enrichment analysis on ethnicity-predictive HM450K sites.** PlaNET's CpG sites used to predict ethnicity was tested for enrichment with respect to a chromosomal location, and b relation to CpG islands.

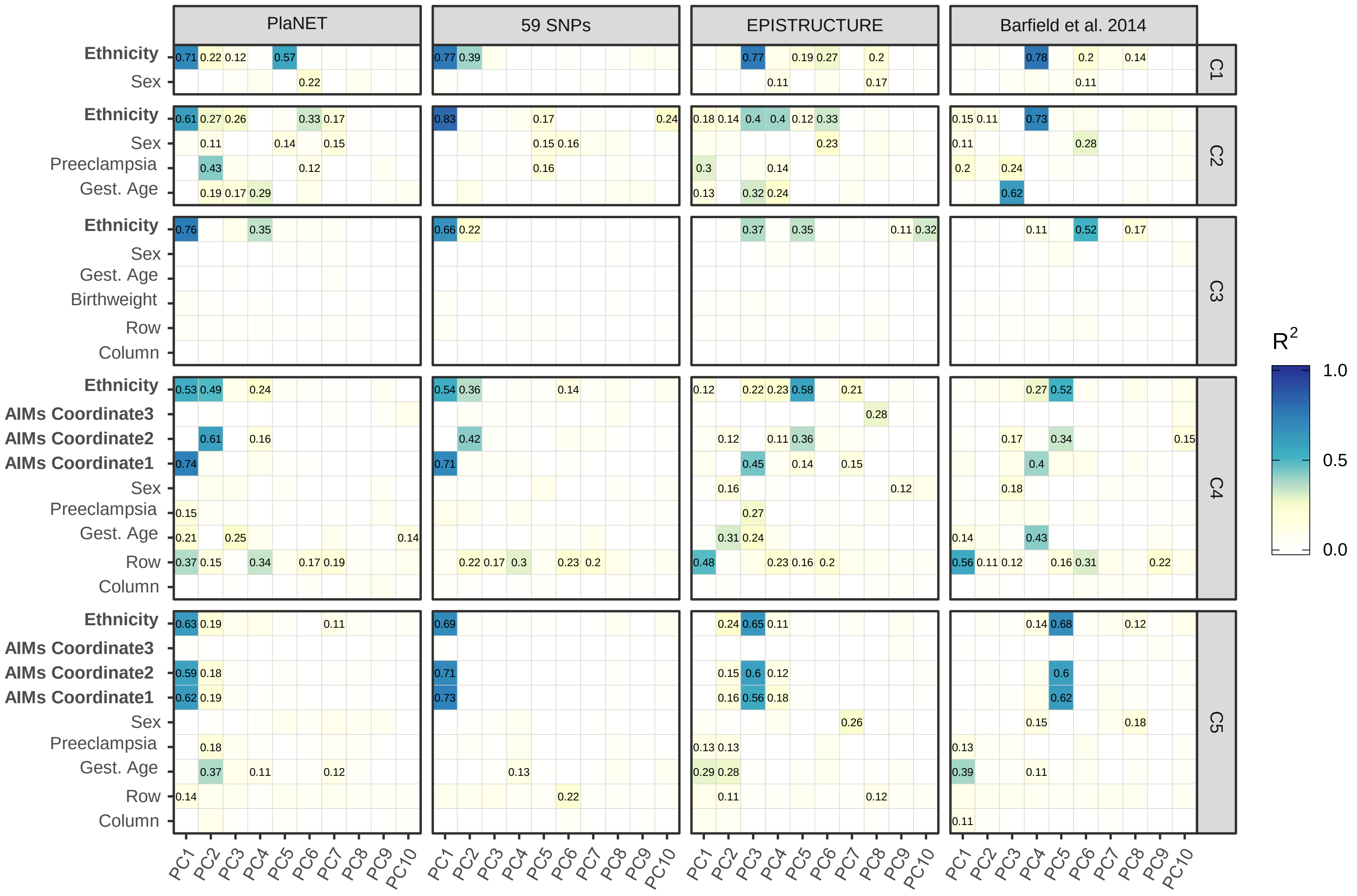


**Figure S6. Association of population structure PCs with technical and biological variables.** PCs were computed on ethnicity-predictive sites, EPISTRUCTURE, Barfield's method and the 59 SNP probes. Each PC was tested for their association with various cohort-specific technical and biological variables. For a given cohort (e.g. C1), ethnicity predictive sites from a classifier trained on all other cohorts (e.g. C2-C5) was used to avoid bias.

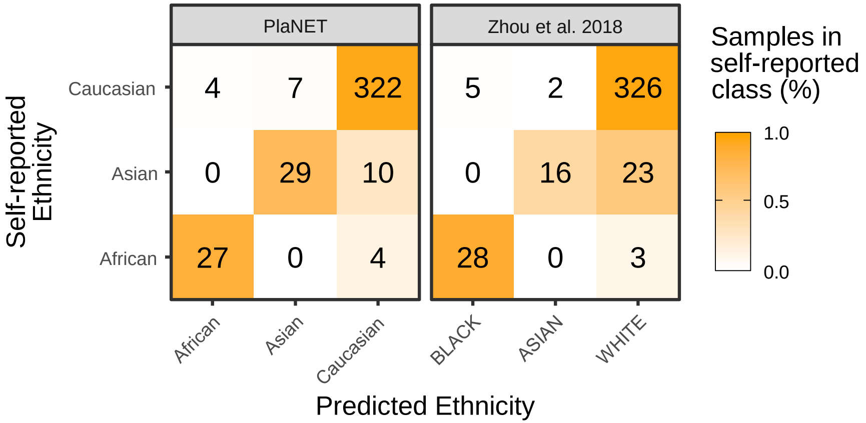


**Figure S7. PlaNET vs Zhou et al. 2017 snp-based classifier.** PlaNET's ethnicity classification performance was compared to Zhou et al. 2018 [76] SNP-based ethnicity classifier in cohorts C3, C4, and C5.

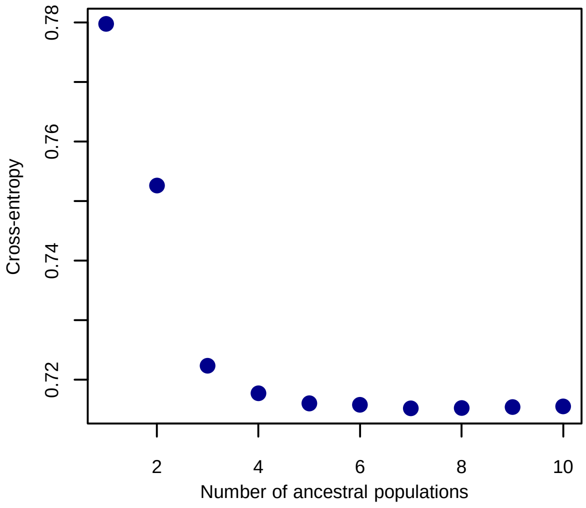


**Figure S8. Estimating k number of ancestral populations using in genetic admixture inference program LEA.** The cross-entropy criterion was used to determine the number of ancestral populations for estimating genetic ancestry coefficients. The number of ancestral populations was chosen at the point k = 3, when the cross-entropy criterion decreases signficantly less with each integer-increase in k.

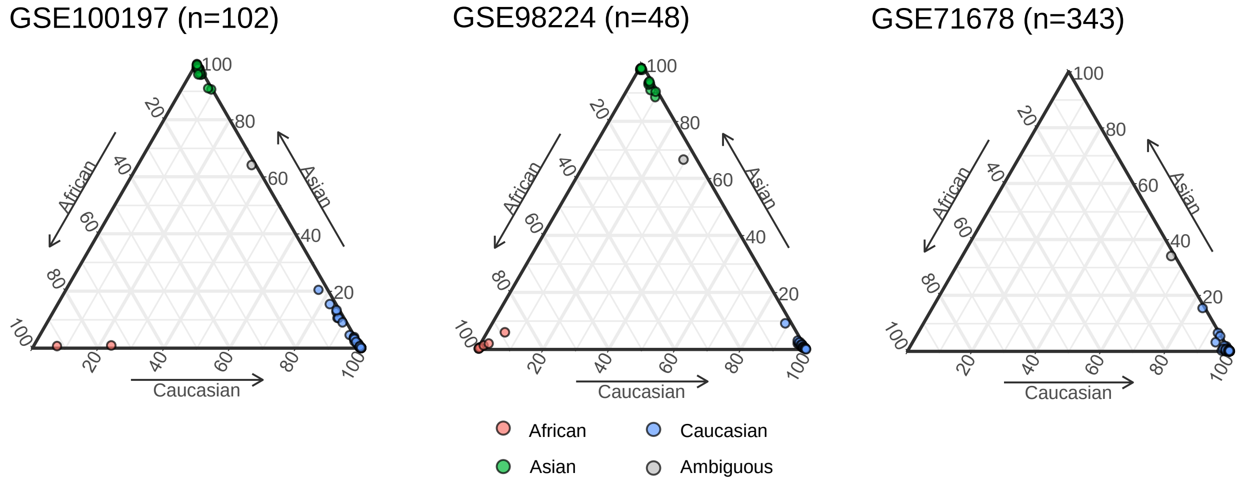


**Figure S9. Application of PlaNET to placental EWAS.** Samples from three independent cohorts are plotted along three axes by their probability of belonging to each ethnicity class and colored by their final ethnicity classification determined by PlaNET.
